# Early 2022 breakthrough infection sera from India target the conserved cryptic class 5 epitope to counteract immune escape by SARS-CoV-2 variants

**DOI:** 10.1101/2024.07.08.602485

**Authors:** Indrani Das Jana, Kawkab Kanjo, Subhanita Roy, Munmun Bhasin, Shatarupa Bhattacharya, Indranath Banerjee, Subhasis Jana, Arjun Chatterjee, Alok Chakraborty, Suman Chakraborty, Budhaditya Mukherjee, Raghavan Varadarajan, Arindam Mondal

## Abstract

During the COVID-19 pandemic, the vast majority of epitope mapping studies have focused on sera from mRNA-vaccinated populations from high-income countries. In contrast, here we report an analysis of 164 serum samples isolated from breakthrough infection patients in India during early 2022 who received two doses of the ChAdOx viral vector vaccine. Sera were screened for neutralization breadth against wildtype, Kappa, Delta, and Omicron BA.1 viruses. Three sera with the highest neutralization breadth and potency were selected for epitope mapping using charged scanning mutagenesis coupled to yeast surface display and Next Generation Sequencing. All of these sera primarily targeted the recently identified class 5 cryptic epitope along with the class 1 and class 4 epitopes, targeted to a lesser extent. The class 5 epitope is completely conserved across all SARS-CoV-2 variants and for the majority of the Sarbecoviruses. In line with these observations, a major fraction of the serum samples, including the selected three, show broad neutralizing activity against recent Omicron subvariants including XBB.1.5. This is in contrast with the results obtained with the sera from individuals receiving multiple doses of original and updated mRNA vaccines, where impaired neutralization of XBB and later variant of concerns was observed. Our study demonstrates that two doses of the ChAdOx vaccine in a highly exposed population was sufficient to drive substantial neutralization breadth against emerging and upcoming variants of concern, thus paves the path towards the development of future vaccine candidates.

**Importance:** Worldwide implementation of COVID19 vaccine and parallel emergence of newer SARS-CoV-2 variants has shaped the humoral immune response in a population specific manner. While characterization of this immune response is important for monitoring the disease progression at population level, it is also imperative for the development of effective countermeasures in the form of novel vaccines and therapeutics. India has implemented the world’s second largest COVID19 vaccination drive and also encountered large number of post vaccination “breakthrough” infections. From a cohort of breakthrough infection patients, we identified individuals whose sera showed broadly cross-reactive immunity against different SARS-CoV-2 variants. Interestingly, these sera primarily target a novel cryptic epitope which was not identified in previous population level studies conducted in western countries. This rare cryptic epitope remains conserved across all SARS-CoV-2 variants including the recently emerged ones and also for the SARS-like coronaviruses that may cause future outbreaks, thus representing a potential target for future vaccines.

## Introduction

Severe Acute Respiratory Syndrome Coronavirus 2 (SARS-CoV-2) has been circulating in the human population for over four years and rapidly evolving, causing the emergence of more than 50 different variants to date (1). Multiple waves of infections were caused by the new variants despite of massive vaccination programs, and their emergence was largely driven by the host immunity generated through natural infections and vaccination (2). Detailed molecular-level understanding of the humoral immunity conferred by different vaccination regimens coupled with ongoing viral exposures is important for the development of effective countermeasures against the current and future variants of SARS-CoV-2 and other Sarbecoviruses that may emerge in future.

More than 5.55 billion people, 72.3% of the world population, have received a single dose of COVID-19 vaccine (3), though in Africa, this number is less than 40% (4). India ranks second highest after China in the number of COVID-19 vaccine doses administered (2.2 billion doses) (5). The vaccination drive started in India on 16^th^ January 2021, at the end of the first wave of infections caused mainly by the circulating B.1.1.7 (Alpha) and B.1.135 (Beta) variants. While the vaccination program was ongoing, the highly infectious B.1.617.2 (Delta) variant emerged and caused the second wave of the pandemic (March to October 2021), which resulted in the highest number of daily infections with high disease severity and mortality rate (6, 7). By January 2022, more than 77% of the Indian population were immunized; the new highly transmissible and heavily mutated B.1.1.529 (Omicron) variant emerged and led to the third wave of infection but was associated with milder symptoms and lower fatality rates (8). A significant fraction of the vaccinated population in India got infected during the second and third waves of the pandemic, leading to the large number of “breakthrough infections” (9). These breakthrough infections shaped the transmission dynamics and contributed to the overall evolution of the virus (10). After 2021, there was little vaccine uptake in the country and the percentage of the population that received a third (booster) dose was only about 10%.

The SARS-CoV-2 spike protein (S) is the main surface glycoprotein that mediates the interaction with the host angiotensin-converting enzyme 2 (ACE2) receptor and subsequent fusion of the viral envelope with the host cell membrane that marks the initiation of the infectious cycle (11, 12). The S protein is the primary target for neutralizing antibodies elicited through natural infection as well as vaccination (13–15). The spike constitutes a trimer where individual protomers are comprised of two subunits, S1 and S2 (16). S1 harbours the N-terminal domain (NTD), the Receptor Binding Domain (RBD), and the S2 houses the fusion peptide, the heptad repeats 1 and 2 (HR1, HR2), the transmembrane domain (TM) and the C-terminal domain (CD). The majority of human neutralizing antibodies, isolated from convalescent or vaccinated sera, target conformational epitopes in the RBD Receptor Binding Motif (RBM) or in the conserved core of RBD (17–19). In the context of the spike trimer, the individual RBDs can be either in the up or down conformation, with only the former able to bind the ACE2 receptor.

The epitopes targeted by neutralizing antibodies can be classified into four major classes (Class 1-4) (20). Class 1 and class 2 epitopes overlap with the ACE2 binding site, and antibodies targeting these sites competitively inhibit receptor binding. While class 1 epitopes are accessible when RBD is present in the up conformation, class 2 epitopes are accessible in both up and down conformations (21). For certain class 2 epitope targeting antibodies (C144), the long complementary determining region (CDR) of the heavy chain acts as a bridge between adjacent RBDs from neighbouring protomers in the down conformation, thus locking the S trimer into the prefusion closed conformation; an unusual mode of neutralization (21). In general, the antibodies targeting class 1 and class 2 epitopes have strong neutralizing activity. However, different SARS-CoV-2 variants escape neutralization by these antibodies by acquiring mutations in the RBM region that is part of the class 1 and 2 epitopes, compromising their efficacy. Both class 3 and class 4 antibodies bind to the RBD core region away from the ACE2 binding site. N343 glycan and the surrounding residues are accommodated in class 3 epitopes (22). S309 antibody, a class 3 antibody isolated from a SARS-CoV infected individual, exhibited broad cross-reactivity against SARS-CoV and SARS-CoV-2, although many SARS-CoV-2 VOCs acquired mutations in class 3 epitopes to escape neutralization by these antibodies (21). The cryptic epitope around the RBD base (first described for CR3022 antibody) constitutes the highly conserved class 4 epitope and is accessible only in the RBD up conformation (23–26). Despite their rare prevalence in convalescent sera, antibodies targeting class 4 epitopes are suggested to protect against emerging VOCs and other sarbecoviruses. Additionally, a highly conserved class 5 epitope has been recently identified and characterized as a target for broadly neutralizing antibodies (27, 28).

Mapping monoclonal antibody epitopes can inform about viral evolution and aid in the development of epitope-based vaccines and therapeutics. However, mapping conformational epitopes targeted by polyclonal sera remains challenging. Mapping these epitopes can elucidate how escape mutations in the viral antigen impact the neutralization potency of the human polyclonal sera harbouring a heterogeneous mixture of antibodies. In the present study, we have conducted a comprehensive analysis of human sera collected from individuals vaccinated with two doses of the ChAdOx1 nCoV-19 viral vectored vaccine and subsequently had undergone breakthrough infections. Neutralization efficacies of 164 sera samples were screened against SARS-CoV-2 pseudoviruses representing the parental strain WA.1 or VOCs B.1.617.1, B.1.617.2 and B.1.1.529. Nine individuals whose serum samples exhibited broader neutralization were subjected to longitudinal sampling and three of them were selected for epitope mapping due to their high neutralization breadth and potency. Utilizing charge scanning mutagenesis (29, 30), a library of individual charged substitutions at all the solvent-exposed positions of the wild-type RBD was generated and displayed on the yeast surfaces to map the epitopes targeted by the polyclonal sera. A high fraction of antibodies targeting either class 1 epitopes or the rare cryptic class 5 epitope, also denoted as RBD8 (31), were observed in all three sera tested. The class 5/ RBD8 epitope residues identified in this study are highly conserved across all SARS-CoV-2 variants, consistent with the observed neutralizing potency of the selected serum samples against heavily mutated Omicron BA.1, BA.5, and the recent XBB.1.5 variant. Our findings illustrate how a combination of vaccination and breakthrough infections in a highly exposed population can elicit diverse antibody classes that confer potent neutralizing activity with broad cross-reactivity. This in turn provides insights into why deaths and hospitalizations have remained low post-2021 in virtually all low and middle-income countries (LMICs) even in the absence of variant updated vaccine boosters.

## Results

### Study cohort

In contrast to vaccination alone, hybrid immunity resulting from SARS-CoV-2 breakthrough infections has been shown to boost neutralizing antibody titers while increasing their breadth at the same time (32, 33). We selected individuals who received two doses of the ChAdOx1 nCoV-19 viral vector vaccine between April and November 2021 and then were infected between November 2021 and January 2022, which marks the transition between the second and third waves of the pandemic in India, caused by the B.1.617.2 (Delta) and B.1.1.529 (Omicron) variants, respectively (34) (Fig 1A, B). We hypothesized that such individuals would have developed broadly cross-reactive neutralizing antibodies that are effective against different variants of concern (32, 33). Blood samples, along with questionnaires about gender, age and symptoms, were collected from 164 selected individuals who were voluntarily recruited by a separate consent for this study. All participants were confirmed positive by RT-PCR for SARS-CoV-2 infection during the abovementioned time frame, which is within 2-4 months post-second dose of vaccination. The serum samples were collected from the individuals within 4-6 weeks of their breakthrough infection (Fig 1A, B). The study cohort consisted of 90 males and 74 females; the median age was 36 and 34 years, respectively (Fig 1C). Most volunteers showed mild symptoms like fever, head and body ache, cough, chest congestion and loss of smell with no cases of hospitalizations. Only 5.84% (10 individuals) of the individuals showed severe to critical illness with declining oxygen saturation and breathlessness along with abovementioned symptoms.

**FIG 1.**
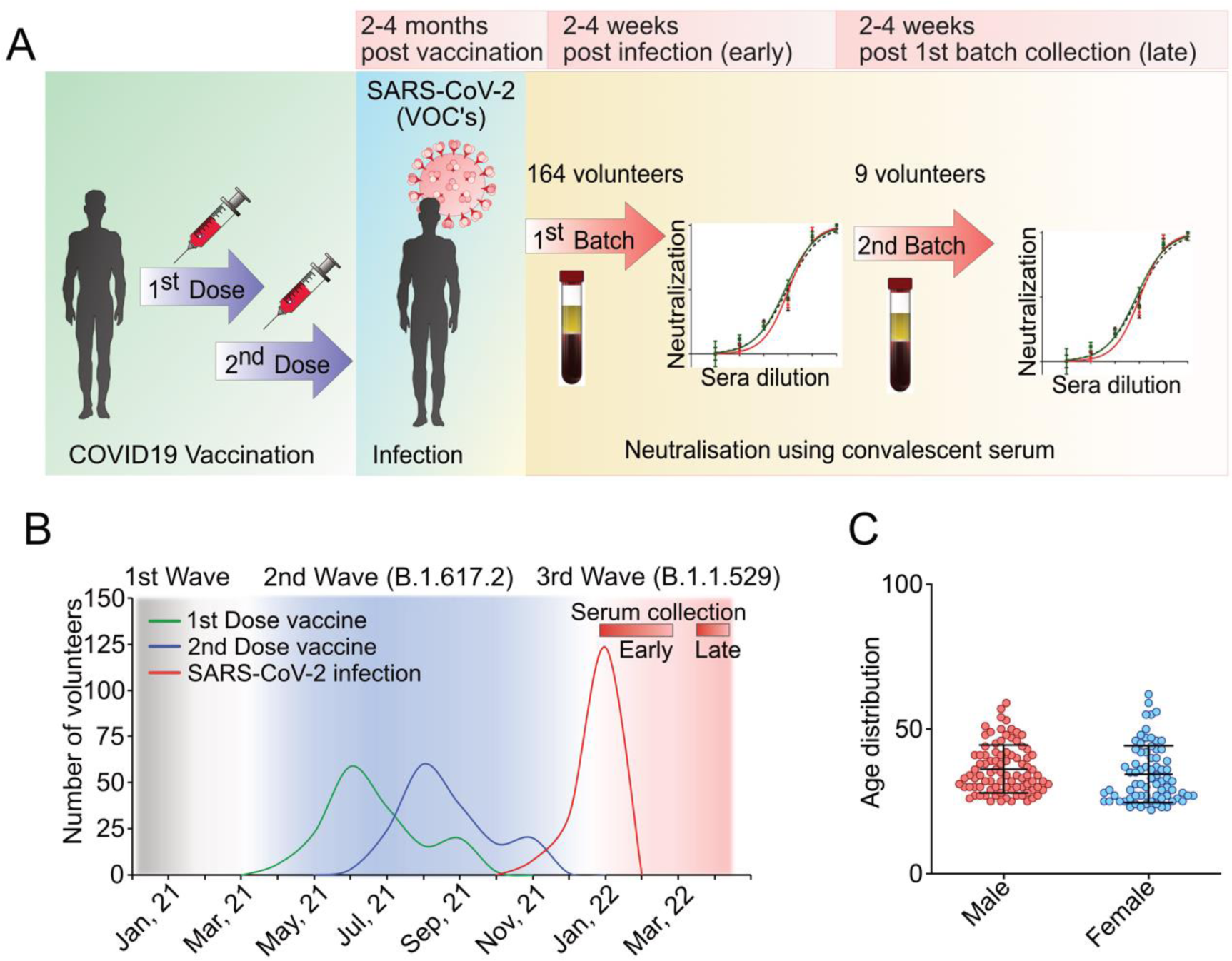
Study cohort and timeline. (A) Schematic representation of study design. (B) Distribution of the study cohort as a function of the timeline of vaccination and breakthrough infection in the context of the second and third wave of pandemics caused by different VOCs of SARS-CoV-2. Red bars indicate the time points of serum sample collection for the entire cohort (early) and subsequently for the selected individuals (late). (C) Age distribution of male and female volunteers included in the study cohort.

### Neutralization profile of the polyclonal serum samples against SARS-CoV-2 variants

We employed a high throughput neutralization assay based on reporter lentiviruses pseudotyped with spike proteins from SARS-CoV-2 parental WA.1, B.1.617.1 (Kappa), B.1.617.2 (Delta) and B.1.1.529 (Omicron) strains (35). For human convalescent sera, such pseudoviruses show neutralization titers comparable to the live SARS-CoV-2, thus can serve as an effective proxy for the live virus neutralization assay (36). Pseudoviruses were incubated with dilutions of serum samples ranging from 10^-2^ to 10^-4^ before infecting HEK-293T cells overexpressing ACE2. The reporter signal was measured as a proxy of the infectivity, which was the highest for the mock-treated cells and the lowest for a polyclonal antibody preparation that binds the spike protein and has high neutralization activity (Novus Biologicals, NB100-56578). The neutralization efficacies of 164 serum samples (all three dilutions) against WA.1 and different VOCs was represented as a heatmap with colours ranging from red (high neutralization) to green (low neutralization) (Fig 2A). Overall, the serum samples from this study exhibited maximum neutralizing activity towards the Delta variant followed by the Kappa variant with a geometric mean (GM) of 93% and 90%, respectively (Fig 2B). Reduced neutralizing activity was observed against the parental WA.1 strain (GM-73%), which was further lowered against the Omicron variant (GM-46%) (Fig 2B). These results suggest that the majority of the individuals in the study cohort had breakthrough infections caused by the Delta variant. Most of the serum samples neutralized Delta and Kappa variants to similar extents, which was better than WA.1 (Fig 2A, B), likely due to the high similarity between the spikes of both variants (37). Interestingly, certain sera (BCRTH 29, 30, 33, 35, 45, 46) showed potent neutralization against WA.1 but failed to neutralize other tested VOCs, especially at higher dilutions (Fig 2A). This suggests that for certain individuals, vaccination and subsequent infections have induced a strong immune response towards the parental strain but failed to generate antibodies that are broadly cross reactive against the VOCs, as also reported previously (38). A few serum samples showed high neutralization activity against the Omicron variant (Fig 2A, B). Limited neutralization activity against the Omicron variant results from the high number of mutations conferring extensive immune escape from vaccine and infection sera (39).

**FIG 2.**
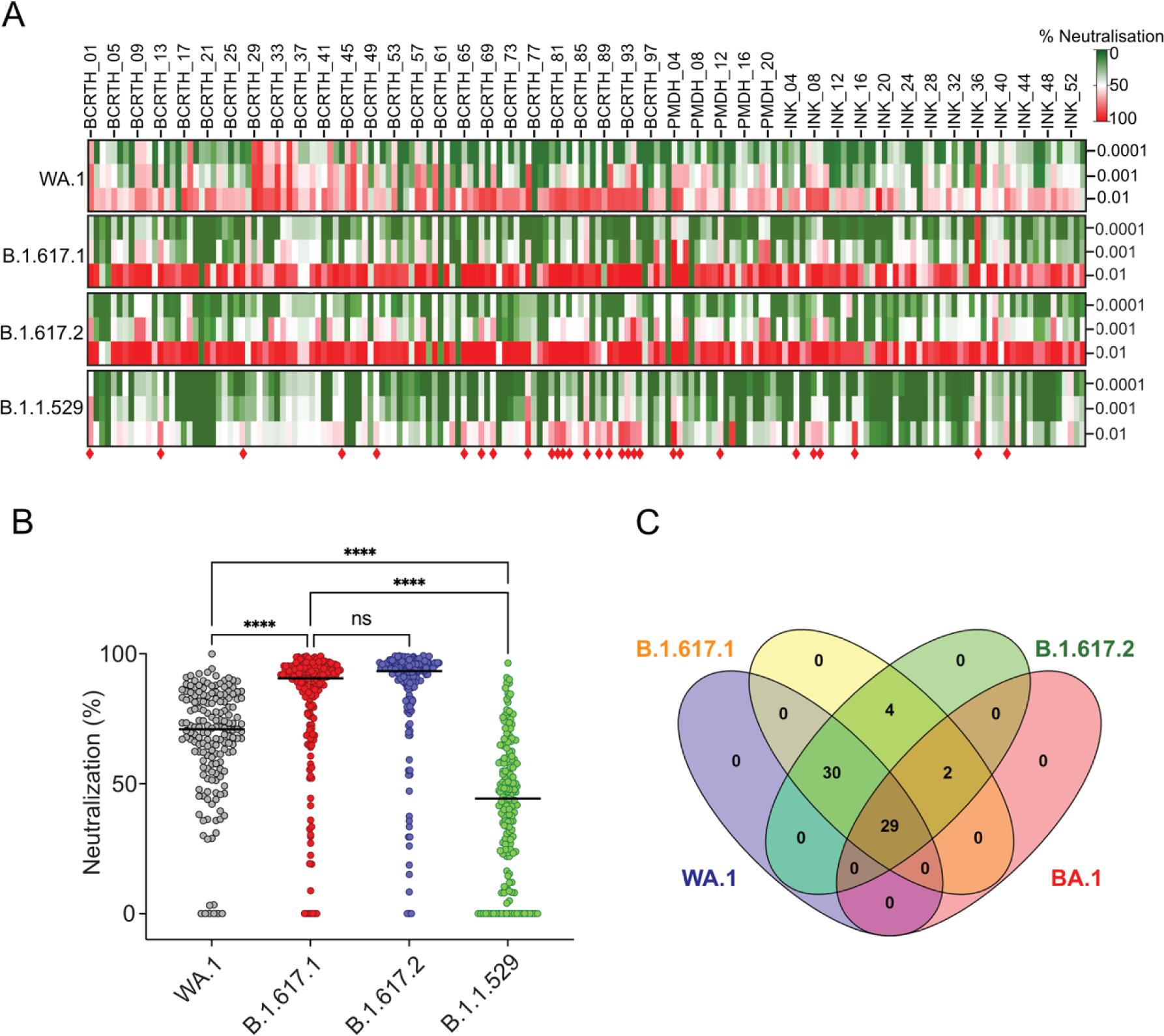
Neutralizing activity of the polyclonal serum samples isolated from the study cohort. (A) Heatmap of neutralizing activity of the 164 serum samples (at dilutions; 0.01, 0.001, 0.001) against the parental strain and different VOCs of SARS-CoV-2 with green to red representing the lowest to highest extent of neutralization (%). (B) Comparison of the relative neuralization efficacies of the serum samples (at 0.01 dilution) against parental strain and different VOCs. (C) Venn diagram showing cross-reactivity of the selected serum samples displaying more than 65% neutralization (at 0.01 dilution) towards at least one of the pseudoviral variants tested in this study. A total of 29 serum samples with >65% cross-reactivity towards all the VOCs tested in this study were designated as broadly cross-reactive sera (red diamonds in A). Each image shown here are a representative of three independent experiments performed in triplicates.

### The breadth and cross-reactivity of the selected polyclonal serum

In the initial screening, 29 out of the 164 breakthrough infection sera (17%) showed broadly cross-reactive neutralizing activity against WA.1, Kappa, Delta and Omicron variants at 10^-2^ dilution (Fig 2C & Fig 2A, red diamonds); hence, were considered as broadly neutralizing serum samples. We tracked these 29 individuals, and selected 9 volunteers who had not been infected and had not received any vaccine booster post the first round of blood collection. Subsequently, second set of serum samples (designated “late”) were collected from the 9 selected individuals with proper consent, 4-6 weeks post the first set (designated “early”) (Fig 1A, B), and both early and late sets of serum samples were subjected to further analysis.

Neutralization efficacies of the selected serum samples were determined against WA.1, Delta and the Omicron variants using the pseudovirus neutralization assay (Fig S1, A, B) and neutralization titers were represented in the form of IC_50_ (Fig 3A, B). Amongst the nine serum samples collected at the early time point, four showed the highest IC_50_ values for Delta, three for the parental WA.1 strain and two for the Omicron; a trend that matched with the overall neutralizing activity of the larger cohort of 164 samples (Delta> WA.1> Omicron; Fig 2B). Interestingly, this was not the case for the serum samples collected from the same volunteers at the later time point (Fig 3A, B). A moderate to sharp decline in neutralizing activity (2-18 fold change in IC_50_) was observed for most serum samples between early and late time points (Fig 3 C, D). Interestingly, a few sera had an increase or maintained similar variant-specific neutralization titers at both time points. Compared to the early time point neutralization profile, BCRTH01 and INK41 showed a significant decline in neutralization titer against WA.1 but a slight increase in neutralization titer against the Omicron variant, at the late time point (Fig 3 C, D). This may reflect the higher durability of the variant specific (Omicron or Delta) antibodies generated by the breakthrough infections with respect to the parental strain (WA.1) specific antibodies elicited through vaccination. Interestingly, BCRTH01, INK09, INK36 and INK41 sera maintained high neutralization titers against all the pseudoviruses tested and at both time points, with INK36 being the most potent neutralizing sera (Fig 3C, 4C and Fig 5B).

**FIG 3.**
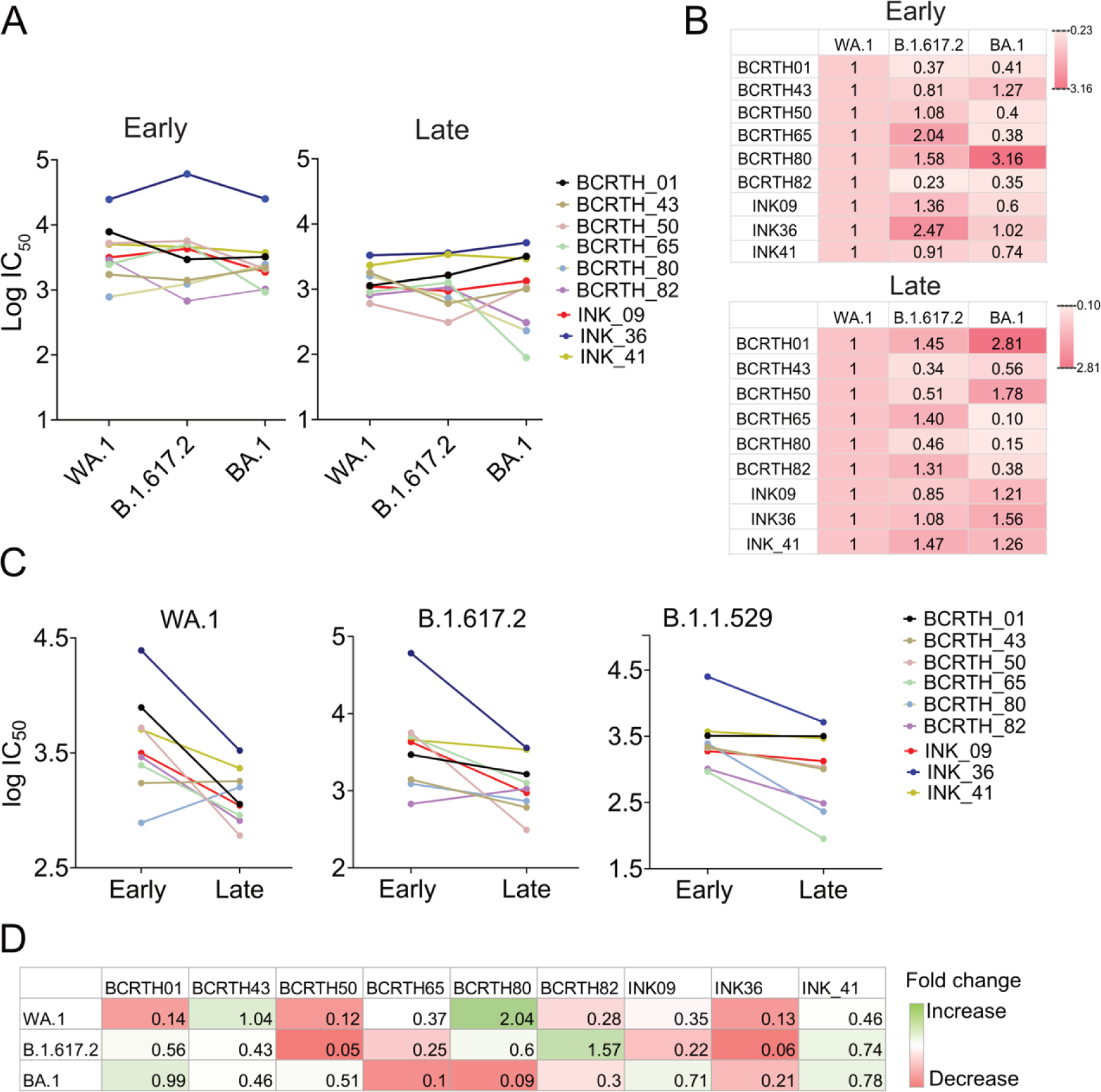
Comparative neutralizing activities of the selected serum samples collected at early and late time points. (A) Comparative analysis of the neutralization titers (IC_50_) of serum samples isolated from the selected 9 volunteers during 1^st^ (early) and second (late) batches of sample collection, against WA.1, B.1.617.2 and BA.1. (B) Heatmap showing fold change in the IC_50_ values for B.1.617.2 and BA.1 with respect to the parental WA.1 strain with red to white indicating higher to lower neutralization titers. (C) Comparative analysis of the pseudovirus neutralization titers of the serum samples isolated at early and late time points. (D) Heatmap representing variant-specific fold change in IC_50_ values. All data shown are averages of the results of at least three independent experiments ± SD.

**FIG 4.**
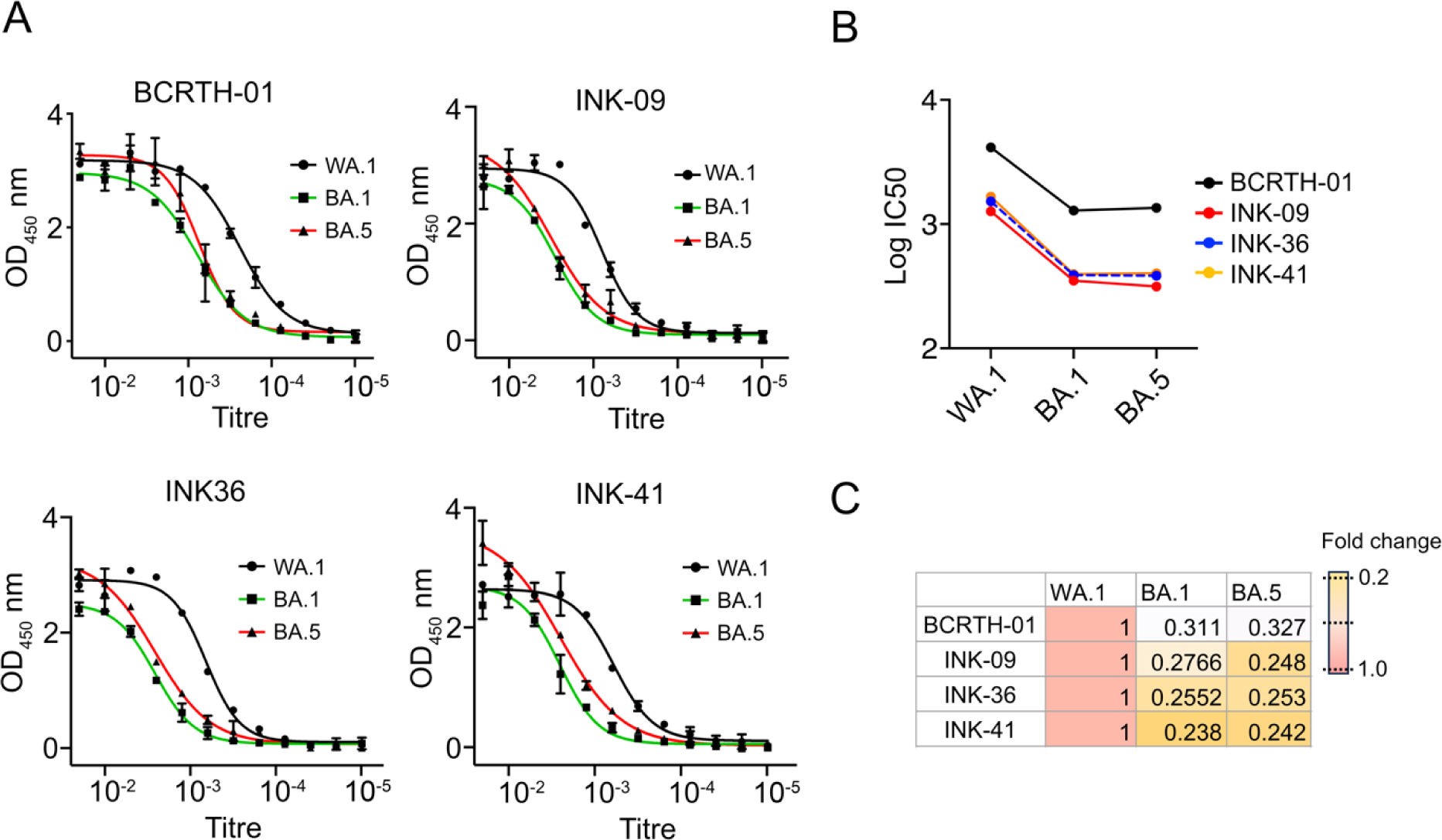
**The serum samples show high binding affinity towards spike RBD**. (A) ELISA was performed to determine the binding affinity of the polyclonal serum samples towards spike RBD derived from WA.1 and Omicron subvariants BA.1 and BA.5. (B) Comparison of binding affinity of different serum samples (presented in the form of IC50) towards different RBD variants. (C) Fold change in IC50 values for the BA.1 and BA.5 variants with respect to the WA.1. Each data shown is a representative of at least three experiments performed independently.

**FIG 5.**
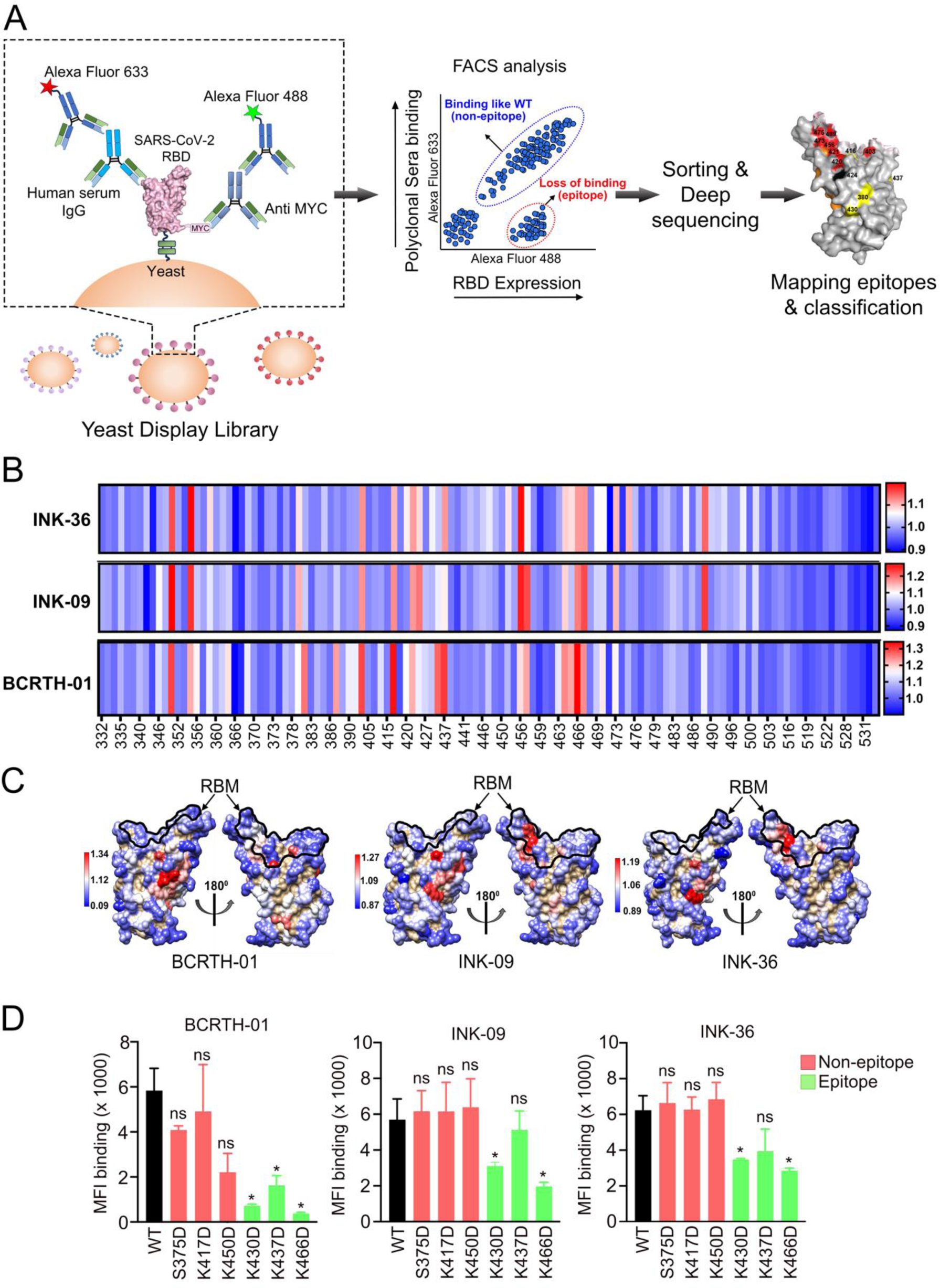
Identification and validation of RBD epitope residues through yeast display analysis. (A) Schematic representation of the yeast display screening as described in the text. (B) The heat map represents the ratio of MFI-expression to MFI-binding extracted from deep sequencing reads analysis. The higher to lower ratios are represented in a red to blue gradient. The higher the ratio (*Red*), the more sera escape at that position. (C) The MFI-expression/ MFI-binding ratios are mapped on the RBD structure (PDB ID: 6M0J) following the same color scheme. The RBM is demarcated on the structure using a black line. (D) Validation was carried out using FACS by quantifying the binding of WT and the mutant RBDs, harboring mutations at the epitope and non-epitope residues, to BCRTH-01, INK09 and INK36 serum samples. The MFI-binding of WT and mutant RBDs w.r.t to the three serum samples are presented. Each image is a representative of two independent experiments performed in triplicates.

As the majority of the neutralizing antibodies target the receptor binding domain (RBD) of the spike protein (17–19), we sought to test the binding activity of the four serum samples (BCRTH01, INK09, INK36 and INK41) to the spike RBD. Enzyme-linked immunosorbent assay (ELISA) was performed with recombinant purified spike RBD derived from the WA.1, Omicron BA.1 and BA.5, which are used as coating antigens. As expected, all four serum samples bound well to the RBD, with the BCRTH-01 sera showing highest binding affinity for all three RBD variants. This suggests high abundance of RBD-targeted antibodies in the selected convalescent serum (Fig 4 A). In general, the serum samples showed highest IgG titres against WA.1 and marginally lower titres against BA.1 and BA.5 (3– 4.2 fold reduction) (Fig 4B, C). BA.5 RBD exhibited higher sensitivity than BA.1, but lower sensitivity than WA.1 for binding to all four serum samples tested. As all of the four serum samples showed comparable neutralization titers against RBD, we selected three of them, namely BCRTH01, INK09 and INK36, for epitope mapping study.

### Comprehensive mapping of RBD epitopes targeted by the selected human serum

We employed a charged scanning mutagenesis library of SARS-CoV-2 RBD displayed at the yeast surface as a novel approach to identify the epitopes targeted by the antibodies present in the polyclonal serum samples (Fig 5A). WA.1 RBD (332-532) was used as the template for library construction and all RBD residues with solvent accessibility >15% were individually mutated to negatively charged Aspartate, with the exception of negatively charged residues, which were mutated to the positively charged Arginine. The generated RBD library was displayed on the yeast surface and subjected to binding with a saturating concentration of the polyclonal serum sample as determined by sera titration on the yeast surface (Fig S2 A). Binding was detected using fluorescently labelled antibodies, and the yeast cell populations were sorted into different bins through Fluorescence Activated Cell Sorting (FACS) (FigS2, B-D). Yeast populations within these bins were subjected to deep sequencing to identify the mutations responsible for abolishing binding to the serum. The deep sequencing reads were analyzed as described in the methods section. Briefly, the binding and expression signals of each mutant RBD were normalized with respect to the wildtype and the MFI ratio between the normalized Mean Fluorescence Intensities for expression (MFI-expression) and binding (MFI-binding) for each of the RBD variants was calculated.

Epitope mapping was performed with all three of the selected serum samples (BCRTH01, INK09 or INK36) and RBD residues whose mutation resulted in MFI_Ratio_ one standard deviation higher than the mean MFI_Ratio_ were considered epitopes (Fig 5B, Table 1). A high value of MFI_Ratio_ (darker shades of red) indicates that the charged substitution significantly perturbed the binding of the serum samples with minimum or no effect on expression. The MFI_Ratio_ of the identified epitope residues were mapped on the RBD structure (Fig 5C). Overall, the identified epitopes (highlighted in lighter to dark shades of red) for all three serum samples were clustered in two major groups; (i) epitopes that overlap with the ACE2 binding motif (RBM-designated by black line on the RBD structure, Fig 5C) and (ii) epitopes that are away from the RBM. Interestingly, for BCRTH01 serum, an enrichment in non-RBM over RBM epitope residues was observed. However, this was not the case for INK36 or INK09, where epitope residues across different RBD regions were enriched to similar levels (Fig 5C).

**Table 1.** MFI ratios for the potential epitope residues.

| SI No. | Residue | MFI-expression/ MFI-binding |  |  |
| --- | --- | --- | --- | --- |
|  |  | BCRTH-01 | INK-09 | INK-36 |
| 1 | 351 | 1.295788 | 1.273718 | 1.157062 |
| 2 | 355 | 1.219729 | 1.239262 | 1.197349 |
| 3 | 356 | 1.17089 | - | - |
| 4 | 359 | - | - | 1.081459 |
| 5 | 364 | 1.182295 | - | - |
| 6 | 380 | - | 1.148814 | 1.077429 |
| 7 | 381 | 1.260786 | - | - |
| 8 | 387 | 1.230637 | - | - |
| 9 | 403 | 1.292267 | 1.172926 | 1.120017 |
| 10 | 416 | 1.329536 | 1.175896 | 1.101252 |
| 11 | 420 | 1.163761 | - | 1.072276 |
| 12 | 421 | 1.218064 | 1.168982 | 1.115263 |
| 13 | 424 | - | 1.192174 | 1.081695 |
| 14 | 430 | 1.270088 | - | - |
| 15 | 437 | 1.301646 | 1.107961 | 1.117455 |
| 16 | 452 | - | - | 1.076026 |
| 17 | 456 | 1.185845 | 1.252503 | 1.192579 |
| 18 | 457 | 1.225249 | 1.222689 | - |
| 19 | 464 | 1.246659 | 1.128143 | 1.087599 |
| 20 | 465 | 1.225858 | - | 1.080284 |
| 21 | 466 | 1.341557 | 1.187018 | 1.116536 |
| 22 | 467 | 1.242733 | 1.221766 | 1.118558 |
| 23 | 470 | 1.178698 | - | 1.063863 |
| 24 | 473 | 1.169721 | 1.145745 | 1.100281 |
| 25 | 475 | - | - | 1.102217 |
| 26 | 489 | - | 1.234014 | 1.144957 |

To this end, we aimed to validate the results of the yeast-display screening and deep sequencing, by individually mutating the epitope and non-epitope residues in RBD and evaluating their impact upon serum binding. Considering that validating all of the library residues would be difficult, we selected three epitope and three non-epitope residues that are identified in the context of all three serum samples (BCRTH-01, INK-09 and INK-36) and reintroduced these mutations into the RBD individually. Subsequently, extent of binding of these mutant RBDs by all three serum samples were evaluated by yeast surface display followed by flow cytometry (Fig S3 A-C). We selected the non-RBM epitope residues (T430, T437 and R466) as they showed high values of MFI_Ratio_ compared to the wildtype. As shown, charged substitution at the putative epitope residues significantly impacted the binding to the serum samples while charged substitution at putative non-epitope residues had no effect upon the binding (Fig 5 D). The impact of mutating the epitope residues was most prominent for BCRTH-01 and higher than INK-09 and INK-36, possibly because of the highest abundance of the RBD specific antibodies in BCRTH-01 serum. This data validates the epitope residues identified through the current study and establishes the robustness of the charged scanning mutagenesis approach for the identification of epitope residues targeted by polyclonal human serum.

### Antibodies in the selected sera target a cryptic epitope that is broadly conserved across different SARS-CoV-2 variants

Previous epitope mapping studies on convalescent polyclonal serum samples showed enrichment of antibodies targeting the class-1, -2 and -3 epitopes and to a lesser extent the class-4 epitope (39–43). Hence, we analysed the different epitopes identified in the present study and compared them to the epitopes reported in the literature. The majority of the neutralizing epitope residues in the RBD are categorized into four major classes, class 1-4 (Fig 6 A and Table 2). Out of the 26 epitopes residues identified, eight were classified as Class 1, two as Class 2, two as Class 3 and five as Class 4. Surprisingly, 11 out of the 26 (42%) epitope residues do not belong to any of these major classes (except for K356 which is shared with class 3 and T470 which is shared with class 2), but instead belong to a rare epitope (44, 45), situated away from the receptor binding motif of RBD (Fig 6A). These residues are part of a recently identified cryptic epitope designated as a class 5 or RBD8 class epitope. (31).

**FIG 6.**
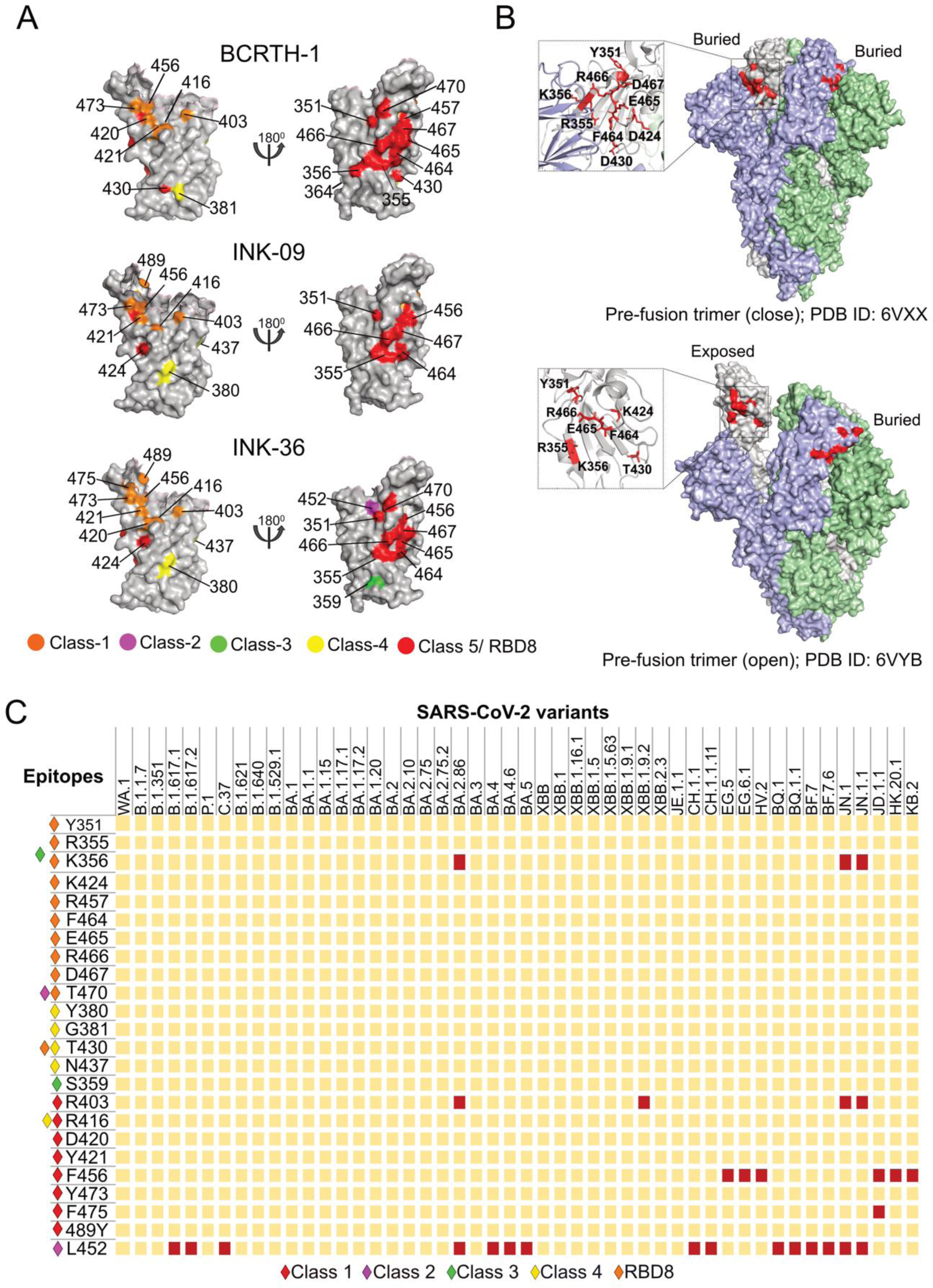
Epitope classification and characterization. (A) Individual epitope residues, as identified for the BCRTH01, INK09 and INK36 serum samples, are mapped on the surface of the RBD (PDB ID: 6M0J) and are color-coded based on the previously published epitope classes. (B) Class 5/ RBD8 epitopes identified in this study are mapped on the pre-fusion trimeric structure of the spike in its closed (PDB ID: 6VXX) and open (PDB ID: 6VYB) conformations. (C) Matrix representing the conservation of identified epitope residues across all major SARS-CoV-2 variants. Mutations of a specific residue in a particular variant are represented in red square. Epitope classes are designated using coloured diamonds.

**Table 2.**
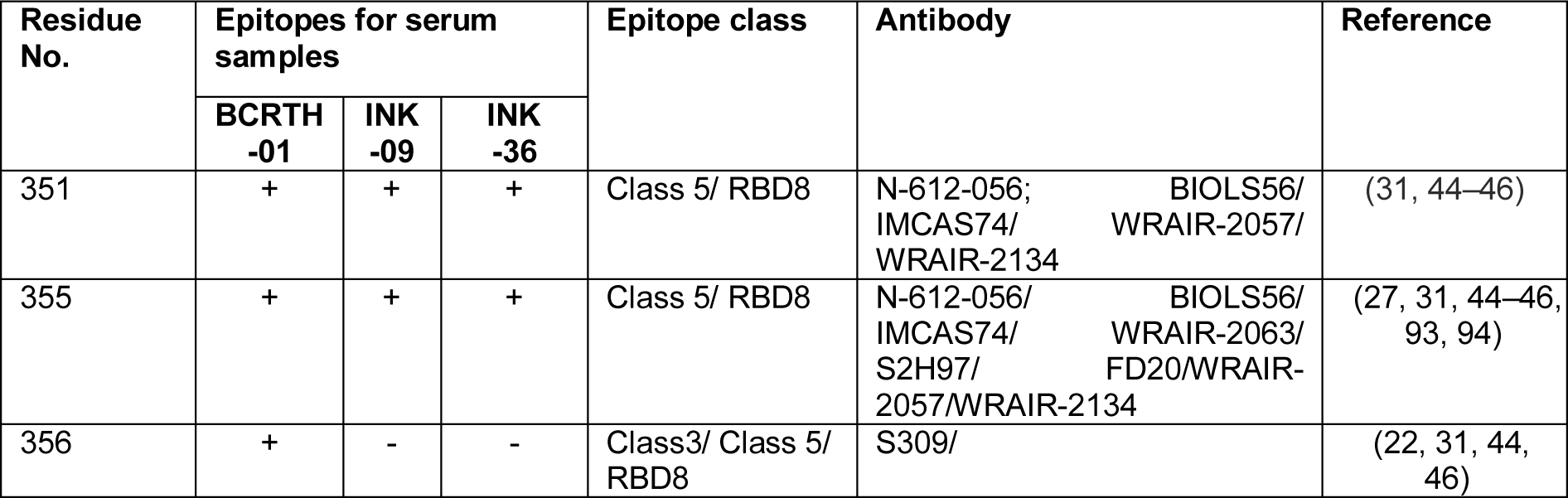
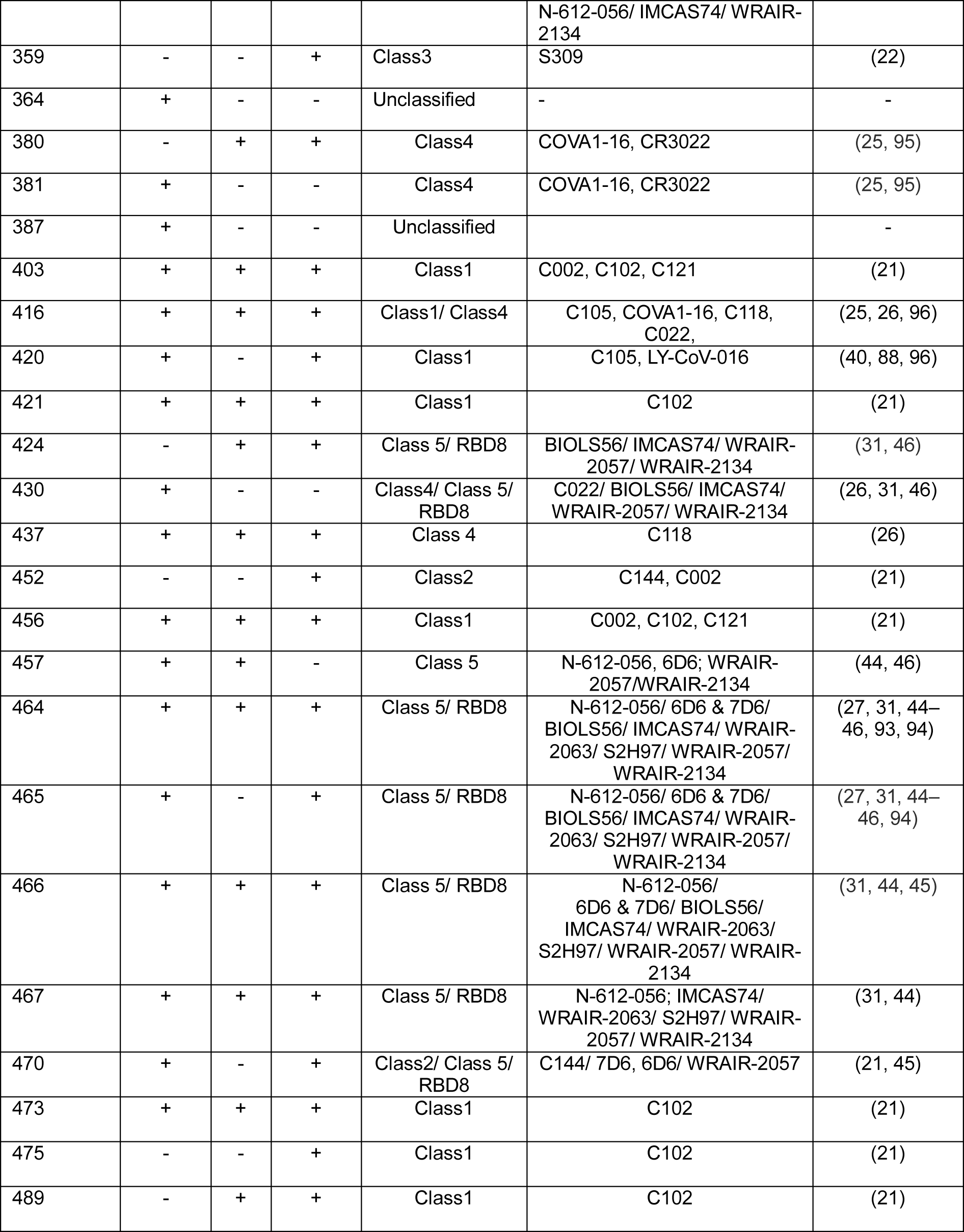
Classification of epitopes.

The class 5 or RBD8 epitope spans diagonally across the base of the RBD (Fig 6A) and is situated on the opposite surface of the ACE2 binding motif, thereby remaining buried and inaccessible in the prefusion closed conformation of the spike trimer (Fig 6B). Hence, binding of class 5 antibodies, such as BIOLS56, IMCAS74, WRAIR-2063/ S2H97/ WRAIR-2057/ WRAIR-2134, to this rare epitope requires extensive opening of the pre-fusion structure, thereby inducing rapid and premature transition to the post-fusion conformation and S1 shedding (27, 31, 46, 47). These antibodies cross-reacted with all SARS-CoV-2 VOCs, including Delta, Omicron BA.4/BA.5, BQ.1.1, and the recent highly immune escaping variant XBB1(31, 46). Supporting this notion, class 5 epitope residues identified in our analysis were highly conserved across all the different SARS-CoV-2 VOCs (Fig 6C) and also across different Sarbecoviruses (Fig S4). All these data together highlights the class 5 antibodies as potential broadly cross reactive universal vaccine candidate against SARS-like betacoronaviruses. Interestingly, the BCRTH01 serum had the highest number (total 10) of class 5 identified epitope residues compared to the INK09 and INK36 (7 and 8 residues, respectively) (Fig 6A, Table 2), which explains its breadth and high cross-reactivity against the different SARS-CoV-2 variants tested. In contrast, INK36 has the highest number of class 1 and class 2 epitopes (8 and 2, respectively) that are targeted by ACE2 competing antibodies, consistent with its highest neutralization activity amongst the three selected serum samples.

To evaluate the effect of charged substitutions on neutralizing activity of the polyclonal sera, we selected four of the identified RBD-epitope residues and individually mutated them to charged substitutions in the WA.1 backbone to generate mutant pseudoviruses. Two mutations, R403D and G416D, are situated within the RBM and are expected to perturb class 1 or class 1/ 4 epitope-specific antibodies, respectively while the other two mutations, T430D and R466D, are situated away from the RBM. T430D is expected to interfere with the binding of class 4 and class 5/ RBD8 epitope-specific antibodies, while R466D is expected to exclusively disturb the latter (Fig 6C, Table 2).

We tested the sensitivity of WT and mutant pseudoviruses to neutralization by BCRTH01, INK09 and INK36 serum (Fig 7A). As expected, all of the mutations reduced the neutralization activity of all three polyclonal serum in comparison to the WT pseudovirus. BCRTH01 showed highest sensitivity towards R466D and R403D mutations with drastic 132-fold and 48.9-fold reduction in the neutralization titer (IC_50_), respectively (Fig 7 B). In contrast, T430D and G416D mutations showed moderate and comparable effects with 3.55- and 4.12-fold reduction in IC_50_ values, respectively. On the other hand, INK36 showed high 35.5-fold and 27.8-fold reduction towards R403D and G416D mutations, moderate 11.5-fold reduction towards R466D mutation and negligible 1.3-fold reduction for T430D mutation respectively. INK09 showed same trend as INK36, but with less prominent reductions towards the abovementioned mutations. It is to be noted that T430 residue has been identified as epitope for BCRTH01, but not for the INK09 or INK36 serum samples and hence its mutations showed negligible reduction towards the neutralization titers of the these two sera. Together, our data suggests that the BCRTH01 sera harbours the highest fraction of class 5/ RBD8 and class 1 antibodies while the INK36 and INK09 possesses higher fraction of class 1 and class 4 antibodies and lower fraction of the class 5/ RBD8 antibodies. This combination of ACE2 competing (class 1) and non-competing (class 5/ RBD8 and class 4) antibodies imparts high neutralization efficacy and broad cross reactivity to all of these serum samples, which render it effective against different variants of SARS-CoV-2, including the currently circulating ones.

**FIG 7.**
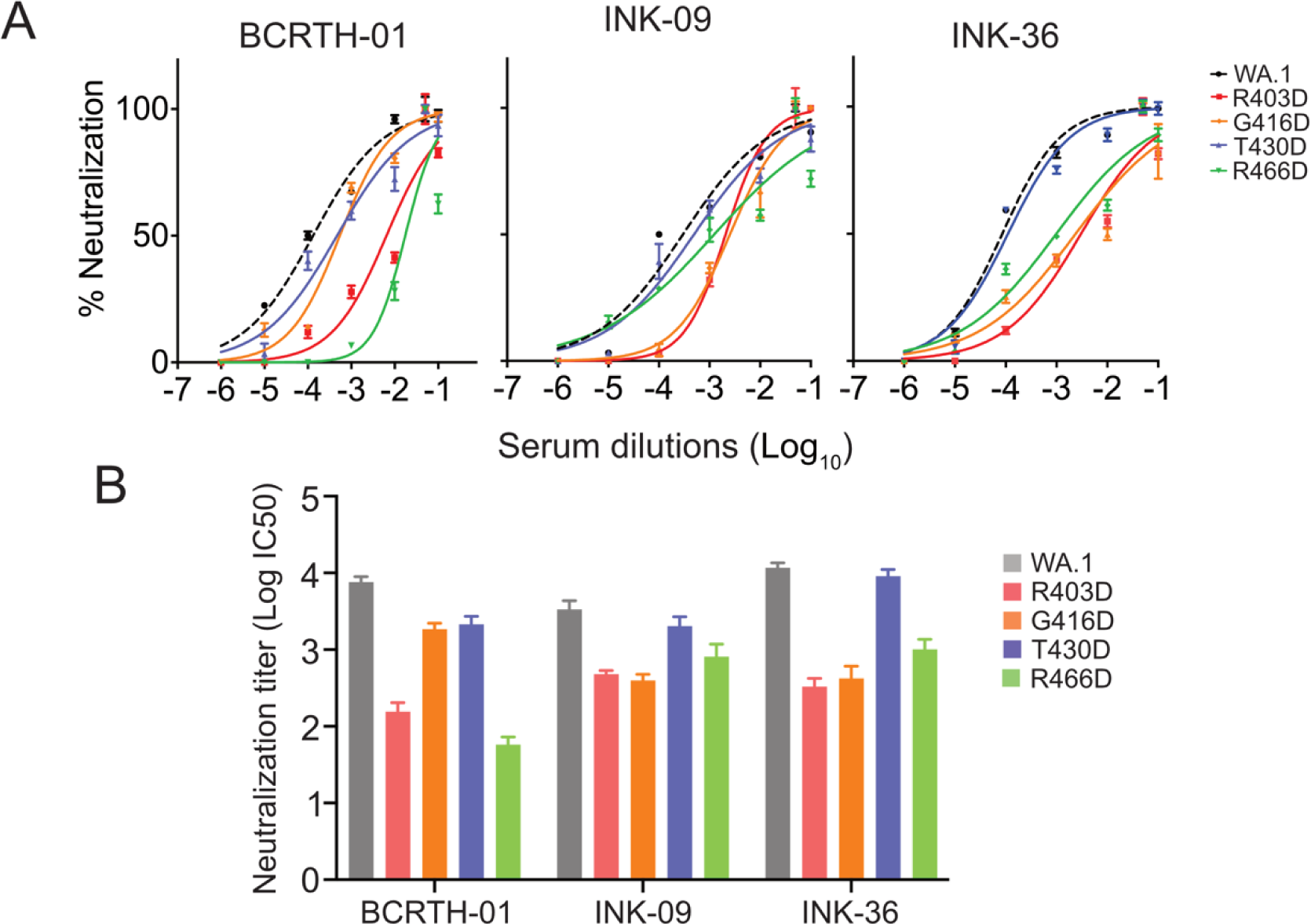
Neutralisation titers (IC50) for WT and mutant pseudoviruses w.r.t selected serum samples. (A) Neutralisation curves for pseudoviruses with WT (WA.1) or mutant spike proteins, harbouring specific mutations at the epitope residues, with serial dilutions of the BCRTH-01, INK-09 and INK-36 sera as indicated. (B) Neutralisation titers (IC50) for WT and mutant pseudoviruses for three sera samples as indicated.

### Pre-XBB serum neutralizes the recent Omicron subvariant XBB.1.5

The heavily mutated Omicron subvariant, XBB and XBB.1, emerged in late 2022 and soon, XBB and its descendants accounted for the majority of infections in the whole world (48, 49). The variant XBB.1.5 harbours a rare mutation, F486P, which demonstrates superior transmissibility and immune escape towards vaccines and monoclonal antibodies, accounting for its rapid spread in more than 100 counties (50–55). Interestingly, the epitopes identified in this study remain completely conserved for XBB.1.5 (Fig 6C), suggesting that the selected serum samples should neutralize this more recent subvariant of SARS-CoV-2. Thus, we generated pseudoviruses harbouring either WA.1, Omicron BA.1 or Omicron XBB.1.5 spike proteins at their surface and performed neutralization assays with the BCRTH01, INK36 and INK09 serum samples which were collected in early 2022, well before the emergence of XBB. Interestingly, all three serum samples showed neutralizing activities against Omicron BA.1 and XBB.1.5 comparable to the parental WA.1 strain (Fig 8A).

**FIG 8.**
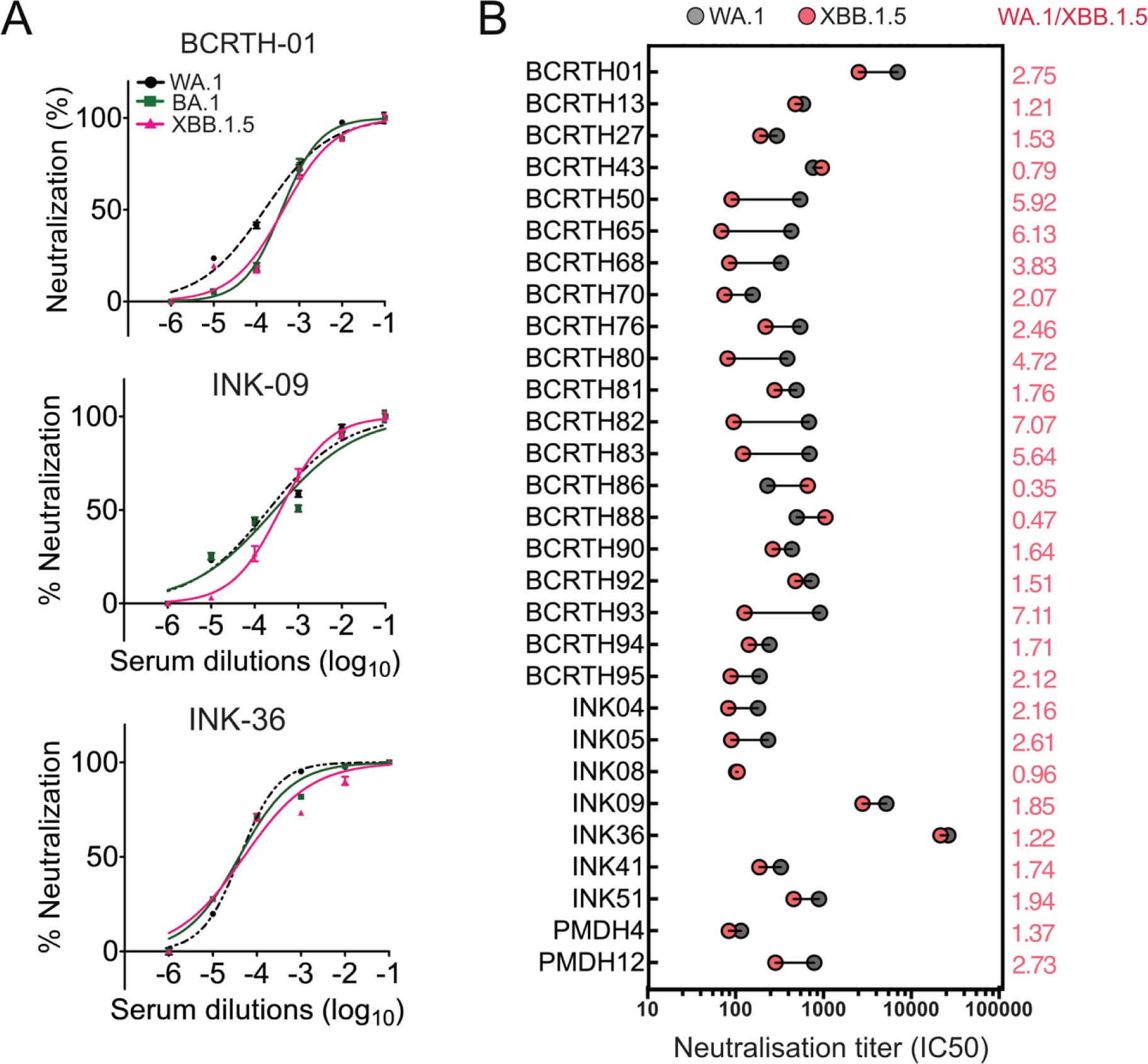
A considerable fraction of the study cohort shows neutralization activity against Omicron XBB.1.5. (A) Neutralisation curves for WA.1, BA. 1 and XBB. 1.5 pseudoviruses with serial dilutions of BCRTH-01, INK-09 and INK 36 sera. (B) Neutralisation titers (IC50) of 29 serum samples (from FIG. 2C) w.r.t WA.1 and XBB.1.5 pseudoviruses are plotted. Fold change of the IC50 for XBB.1.5 w.r.t the WA.1 are marked in salmon red. All data shown are averages of the results of at least three independent experiments ± SD. ∗= P < 0.001,∗∗∗∗ = P < 0.0001 (unpaired Student’s t-test). Images shown are representative of three independent repeats.

It is possible that the XBB.1.5 neutralization activity showed by the three selected serum samples may be an outcome of random stochasticity and do not represent a general trend of the larger study cohort. To investigate this, we evaluated the neutralizing activity of the all 29 serum samples, which showed broader cross-reactivity against all of the tested SARS-Cov-2 variants in the initial screen (Fig 2C). WA.1 or the XBB.1.5 pseudoviruses were neutralised with different concentration of the serum samples (Fig S5) and the respective neutralization titers are plotted in Fig 8B. As evident, all of the serum samples showed comparable neutralizing activity towards WA.1 and XBB.1.5 with a maximum of 7 fold reduction for the later. This data indicates that a significant fraction (17%) of the participating volunteers, who had breakthrough infections following vaccination during the Delta/ Omicron wave (December 2021) of the pandemic, generated humoral immunity that remained effective even against recently emerged subvariants (February-March 2023). This, in turn, establishes the potential importance of the identified epitopes as a target for developing vaccine candidates against future SARS-CoV-2 variants.

## Discussion

Hybrid immunity generated by breakthrough infections have been reported to induce enduring humoral immunity and generate broadly neutralizing antibodies (33, 56, 57) against SARS-CoV-2 variants through several different mechanisms (58). First, vaccination and infection select different B cell clones leading to the generation of broadly cross-reactive neutralizing antibodies (59, 60). Second, the antibodies generated through primary antigen response directly modulate the secondary response against SARS-CoV-2 by blocking or enhancing the naïve B cell recruitment (61). Third, timely spaced repeated antigen contacts through vaccination and subsequent infection promote more extensive germinal centre reactions resulting in high affinity cross-reactive neutralizing antibodies (62, 63). These together explains why breakthrough infections with specific omicron variants activates specific memory B cells that recognizes conserved cross-variant epitopes rather than promoting naïve B cell recruitment (64, 65). In this study we examined the hybrid immunity and characterized the epitopes targeted by the antibodies generated in a small cohort of breakthrough infection patients from India.

Human high-resolution epitope mapping studies following vaccination and/or infection have been largely focused on sera from US or European populations that were primarily immunized with mRNA vaccines (19, 43, 66, 67) while a few studies focused on sera from individuals vaccinated with CoronaVac whole virus inactivated vaccine, with or without breakthrough infection (54, 68, 69). Little is known about the humoral response in LMICs and China, which are home to ∼85% of the world’s population. In the present study, we describe epitope mapping of sera from an Indian population where participating volunteers received two doses of viral vector vaccine and had a “breakthrough infection” within 4-6 weeks post-second dose. The timing of the study was deliberately chosen at the intersection of the second and third waves of the pandemic instigated by the B.1.617.2 (Delta) and B1.1.529 (Omicron) variants, which, in theory, were expected to enhance the breadth and potency of antibodies generated through vaccination (parental strain WA.1) and subsequent infections by different VOCs (33). Around 18% of the serum samples exhibited > 65% neutralizing activity against the parental strain and different VOCs including Kappa, Delta, and Omicron (Fig 2C). Furthermore, via longitudinal sampling, three serum samples were identified that not only maintained their broad neutralizing activity up to 12 weeks post-infection but also demonstrated effectiveness against the recent circulating Omicron subvariant XBB.1.5. Although prior exposure to Omicron spike, including BA.1 bivalent vaccination has been reported to generate XBB.1.5 specific antibodies (70), the neutralizing activity remained low compared to the parental WA.1 or D614G strains (52, 70–72). Several studies have demonstrated a substantial escape of XBB variants from neutralization by convalescent and vaccinated sera, with more than 20-fold reduction in neutralization titres compared to WA.1 (41, 54, 71, 73). Strikingly, 29 out of 164 (18%) broadly cross reactive serum samples (from Fig 2C) showed comparable WA.1 and XBB.1.5 neutralizing titers (≥ 7 fold reduction for XBB.1.5) (Fig 8B), hence representing proof towards the role of hybrid immunity in generating highly potent, durable, and broadly cross-reactive neutralizing antibodies.

We employed an aspartate scanning mutagenesis and yeast surface display coupled with flow cytometry and deep sequencing to map epitope residues which, upon charged substitution, severely impacted the binding of the serum antibodies without affecting the expression. Charge scanning mutagenesis has been previously proven to be effective in identifying residues buried at protein-protein interaction interface (74, 75). While Deep Mutational Scanning has been previously employed to identify serum escape mutations on SARS-CoV-2 RBD (19, 76), the relatively smaller size of the aspartate scanning library, together with the unique barcoding strategy of the individual mutant ORFs employed by us simplify library construction and downstream analysis. Using this method, we identified a total of 26 surface-exposed amino acid residues as potential epitopes targeted by the sera. The majority of these residues have been previously identified as part of epitopes for different classes of RBD neutralizing monoclonal antibodies (77), thus supporting the robustness of the novel screening approach in identifying epitopes targeted by polyclonal sera.

The epitopes identified using the proposed analysis can be broadly categorized into two main groups: RBM epitopes overlapping with the ACE2 binding motif and non-RBM epitopes. The RBM epitopes consist primarily of class 1 and class 2 epitopes which are targeted by antibodies that competitively hinder receptor engagement and virus attachment to host cells. While antibodies targeting these epitopes are responsible for the high neutralizing activity of polyclonal serum used in this study (17, 21, 78–80), higher frequencies of escape mutations in these regions limits their breath and cross-reactivity in the emerging VOCs (25, 26, 47, 81). For example, the R403K mutation, in the class 1 epitope, has been classified as “vaccine escape mutation”, leading to higher infectivity and lower neutralizing activity of vaccine-elicited antibodies against the recent JN.1 subvariant of Omicron (82). Interestingly, the R403D mutation resulted in 48.9 fold reduction in neutralizing activity of the BCRTH01 serum as evidenced in our study. Another identified class 2 epitope residue substitution L452R, was associated with a massive expansion and immune escape of the Delta variant from vaccine-elicited class1 and class 2 antibodies (43) during the second year of the pandemic (83). Similarly, the F456L mutation (class 1 epitope) is associated with heightened ACE2 binding, infectivity, and immune evasion from the XBB.1.5 neutralizing antibodies (84) leading to the global spread of EG.5 (Eris) subvariant (85) in the recent past. The K356T mutation in JN.1, in the class 3 epitope, drives resistance towards Sotrovimab, a class 3 antibody (S309) based antiviral drug (86). In contrast to classes 1-3, the class 4 epitopes targeted by the selected sera described above show high conservation (Fig 6C). These class 4 epitopes are situated in the RBD core region, away from the RBM except for the G416, which resides within RBM. Class 4 antibodies targeting G416 (COVA1-16, C118 and C022) and surrounding residues sterically hinder ACE2 binding (25, 26) thereby resulted in high neutralizing activity.

A major fraction (11/26) of the epitope residues identified in the present study belong to a newly identified epitope (44, 45, 87), rarely seen in the repertoire of antibodies from convalescent sera (40–43). Recently, this epitope has been categorized as class 5 or the RBD8 class of epitopes (27, 31). These antibodies are reported to get enriched by breakthrough infections caused by Omicron subvariants (54). The amino acid residues constituting these epitopes are clustered together at the opposite surface from the ACE2 binding motif of RBD and remain inaccessible in the closed trimeric form of the spike (S) (Fig 6B). Such epitopes can only be accessible through breathing of the prefusion structure of the trimeric S protein (Fig 6B) whereby antibody binding ultimately leads to shedding of the S1 subunit (31). Additionally, it has been proposed that the binding of class 5 antibodies leads to premature, receptor-independent conversion of S from pre-to post-fusion conformation, ultimately blocking viral entry and syncytium formation (88). Interestingly, class 5 epitope residues remain completely conserved across all SARS-CoV-2 variants and for a number of different Sarbecoviruses, indicating the cross-reactivity and increased breadth of class 5 antibodies against these viruses. Consistent with this, the selected serum samples, enriched for RBD8 epitope antibodies, effectively neutralized WA.1, Delta, Kappa, Omicron BA.1 and XBB.1.5.

Lately, antibodies targeting non-RBM epitopes in the spike protein have emerged as prime candidates for developing broad-spectrum vaccines or therapeutics against SARS-CoV-2 variants (28, 89, 90). Some of these antibodies showed cross reactivity against different Sarbecoviruses, thus further extending the prophylactic/therapeutic potential of such antibodies against future emerging SARS-like coronaviruses. However, non-RBM antibodies are limited by their weak neutralization activity. Rao et al., have shown that the RBD8 antibody (BIOLS56) can rescue the immune escape caused by an RBM targeting antibody (IMCAS-L4.65), together ensuring strong neutralizing activity and broad neutralizing efficacy against different Omicron variants (31). In this study, we show that an immune response elicited by hybrid immunity and containing a combination of class-1 (RBM targeting) and class 5/ RBD8 antibodies can achieve neutralization potencies against different SARS-CoV-2 variants including WA.1, Kappa, Delta and multiple Omicron subvariants including the more distant XBB.1.5. Characterizing such combination of different classes of antibodies is important for optimizing novel vaccine recipes against existing and yet to be emerging sarbecovirus. Interestingly, mutation of a single class5/ RBD8 epitope residue (R466D) resulted in 132 fold reduction in the neutralization titer, which is even higher than that observed in the case of mutations of class 1 (R403D) epitope (48.9 fold), suggesting high abundance and comparable neutralization efficacy of these two different classes of antibodies (class 5/ RBD8 and class 1) in the one of the selected serum samples (BCRTH01) used for epitope mapping analysis. This contrasts with the previously conducted epitope mapping studies from the convalescent polyclonal serum samples, which led to the identification of class 1-4 epitope targeting antibodies (40–43).

In conclusion, antibodies targeting different classes of epitopes may synergistically contribute to achieving high neutralizing activity (Class 1+2 epitopes) and broader cross-reactivity (Class 4 and RBD8) against all variants of SARS-CoV-2. Notably, RBD8 constitutes a significant portion of the epitopes identified in all three serum samples, which differs from the typical composition of the convalescent serum samples analysed previously (40–43). In India, after the Delta wave in mid-2021, there was little masking or isolation, and likely continued extensive viral exposure. This, coupled with the relatively young population (median age 28.4 years) may have contributed to the generation of broadly cross reactive antibodies targeting novel cryptic epitope presented here. These findings need to be confirmed in larger epitope mapping studies where the humoral immune response, abundance of specific classes of antibodies should be correlated with the clinical manifestation of the disease. Greater understanding of this adaptive strategy implemented by the host would pave the way towards the development of universal vaccines and therapeutics against newly emerging SARS-CoV-2 variants and future sarbecovirus outbreaks.

## Materials and methods

### Cell line

Human embryonic kidney (HEK) 293T cells (ATCC #CRL-3216) were maintained in Dulbecco’s modified Eagle’s medium (DMEM) supplemented with 10% fetal bovine serum (FBS) along with antibiotics like penicillin, streptomycin and anti mycotic agent (Gibco) at 37°C and 5% CO2.

### Plasmids

Plasmid pHAGE2-EF1aInt-ACE2-WT, expressing human ACE2 gene (BEI#NR-52514) was used for overexpressing ACE2 in HEK 293T cells. Plasmid HDM-IDTSpike-fixK (BEI#NR-52514) was used for expressing the codon-optimized spike protein from SARS-CoV-2, Wuhan-Hu-1 strain (Genbank NC_045512). Plasmids pcDNA3.3-SARS2-B.1.617.1, expressing spike protein of Kappa variant (addgene#172319), pcDNA3.3-SARS2-B.1.617.2 SARS-CoV-2, expressing spike protein of Delta variant (addgene#172320) (91) and SARS-CoV-2 Omicron strain spike gene in Human codon_pcDNA3.1(+) expressing spike protein of Omicron variant (B.1.1.529) (GenScript# MC_0101274) were utilized in the study (35). XBB1.5 spike expression plasmid construct was a kind gift from Dr. Rajesh P. Ringe, Institute of Microbial Technology, Council of Scientific and Industrial Research (CSIR), India.

### Antibodies

NB100-56578 is a polyclonal antibody raised in rabbits against SARS CoV-2 spike protein and was obtained from Novus Biologicals (USA). Serum from the pre-COVID era (Single donor human serum off the clot, ISERS2M-2019-33108-07) was obtained from Innovative Research, Inc.

### Bioinformatics and Structural Analysis

Genome sequences of 47 different SARS-CoV-2 viruses that infect humans and 11 different sarbecovirus spike RBD sequences were obtained from the GISAID database. Multiple sequence alignment was performed in Bioedit software using ClustalW multiple alignment function. Further representation of the spike proteins’ aligned RBD amino acid sequences was performed using JALVIEW or the Multlin sequence alignment tool. Structural alignment was performed in Pymol using specific PDB files of post-fusion spike protein structures of SARS-CoV-2 viruses (PDB id: 6M0J).

### Construction of SARS-CoV-2 S mutant plasmid constructs

Plasmid HDM-IDTSpike-fixK expressing a codon-optimized spike protein from SARS-CoV-2 was subjected to site-directed mutagenesis to generate R403D, G416D, T430D and R466D, respectively, using the QuikChange II Site-directed mutagenesis kit (Agilent) according to the manufacturer’s instructions. Mutations generated in the SARS-CoV-2 S protein were further verified using Sangers sequencing.

### Generation of SARS-CoV-2 S pseudotyped lentiviruses

Generation of SARS-CoV-2 lentivirus particles pseudotyped with either full-length WT, different VoCs or mutation at SARS-CoV-2 spike protein (R403D, G416D, T430D and R466D) were generated by transfecting HEK293T cells with the respective set of plasmids by using the protocol as described previously by Crawford et al., 2020 (92). Briefly, a lentiviral backbone plasmid, pHAGE-CMV-Luc2-IRES-ZsGreen-W (BEI #NR-52516) bearing a CMV promoter to express luciferase, was co-transfected with a set of other lentiviral plasmids like pHDM-Hgpm2, expressing the HIV-1 gag and pol (BEI #NR-52517), pRC-CMV-rev1b, expressing the HIV-1 rev (BEI #NR-52519), pHDM-tat1b, expressing the HIV-1 tat (BEI # NR-52518), and respective different WT or mutant spike protein-expressing plasmids. Post transfection, the SARS CoV-2 pseudoviruses were collected by passing the supernatant through a 0.45 um filter and used freshly.

### Pseudovirus-based Serum Neutralization Assay

Serum samples collected from the volunteers were inactivated at 65^0^C for 30 minutes prior to use. Serially diluted serum samples were incubated with either WT or different VoC/ mutant pseudoviruses at 37^0^C for 1h. Serum-neutralized viral inoculum was then used to infect hACE2-expressing HEK 293T cells at 37°C for 1h in the CO_2_ incubator in the presence of polybrene till 18h. Successful neutralization of the pseudovirus results in a several fold reduction in reporter virus activity, which were measured using a multimodal plate reader (GloMax® Explorer Multimode Microplate Reader, Promega). Polyclonal SARS CoV-2 (NB100-56578) antibody served as a positive control, whereas sera from the pre-COVID era were used as a negative control for the study.

### Charged scanning library construction and yeast surface display

Substitutions to Aspartate and Arginine are chosen for introduction in scanning mutagenesis as analysis of existing DMS data suggests that these are the most disruptive for binding. A charged scanning library of SARS CoV-2 spike RBD was generated using site-directed mutagenesis. Exposed positions with accessibility >15% (total 125 residues) were individually mutated to Aspartate or Arginine (for negatively charged residues), and each mutant was fused by PCR to a unique 6 nucleotide barcode of known sequence to simplify analysis and eliminate the need for long read sequencing. Upon mutagenesis by overlap PCR, all the PCR products from each individual mutated position were pooled and transformed into yeast via two fragment recombination by electroporation.

### Human sera screening

Three human sera were selected for screening based on their high neutralization titres. The sera were first titrated on yeast surface displaying WT-RBD and a saturating concentration was chosen from the titration curve to be used for epitope mapping. The binding of these sera with the aspartate library was analysed by flow cytometry, sorted into multiple gates and then subjected to deep sequencing. Briefly, the pooled library of yeast cells expressing each of the charged mutants was inoculated in 5 ml of SDCAA medium and grown overnight at 30°C, 250 rpm. The next day, a secondary inoculation from the overnight culture of the library was grown at 30°C until an OD600 of 3–4, followed by induction in SGCAA medium at 20°C for 16 hr. A total of 1× 10^7^ cells of this library were washed twice with FACS buffer (1XPBS, pH 7.4 and 0.5% BSA) and then incubated at 4°C for 1 hr with a selected dilution of sera to probe for binding and chicken anti-CMyc antibody (1:300 dilution) as a probe for the expression. The cells were washed twice with FACS buffer and subsequently labelled with Alexa Fluor 633 goat anti-human (1:1200 dilution) (catalogue no. A21091), plus Alexa Fluor 488 goat anti-chicken (1:300 dilution), (catalogue no. A11039) for 20 min at room temperature in the dark. After washing thrice with FACS buffer, cells were resuspended in 1 mL of FACS buffer, subjected to flow cytometry, and sorted on an Aria-III cell sorter (BD Biosciences).

### Deep sequencing and analysis

The sorted yeast cells from each gate were grown separately in liquid SDCAA medium to saturation at 30°C for 24 hr, and the plasmids were extracted using EZ Yeast plasmid miniprep kit (G Biosciences) according to the manufacturer’s instructions and then subjected to PCR. The PCR primers were designed to amplify the barcode region only (150 bp) and in a way such that for both, the forward and reverse end reads, the first 3 bases are NNN followed by 6 bases of unique sequence tag (MID) and the 21 bases of the primer sequence complementary to the gene. Each forward MID sequence represents a particular serum, while each reverse MID sequence represents a specific gate. The PCR for all the gates from three sorted sera samples was carried out using Phusion polymerase for 15 cycles. Following agarose gel electrophoresis and gel band purification, an equal amount (∼100 ng) of PCR product from each sample was pooled and then gel-band purified, followed by sequencing on the NovaSeq platform. The fraction of each mutant (*Xi*) distributed across all the bins was calculated as in (Equation 1) and then the reconstructed MFI for each mutant was calculated by the summation of the product obtained from multiplying the fraction of a particular mutant in a particular gate (*Xi*) with the MFI of that gate that was obtained from the FACS experiment (*Fi*), (Equation 2).

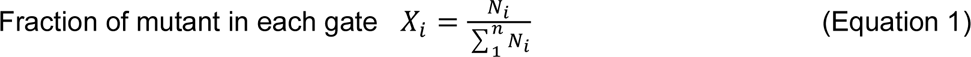

N*i*: Number of reads of a specific mutant in bin *i*.

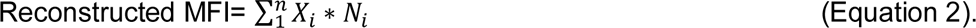

The ratio of MFI expression/MFI binding for all positions was represented by a heat map using GraphPad Prism software. The mutants with Ratio (Normalized MFI of expression: Normalized MFI of binding), one standard deviation higher than the mean MFI were considered as epitope residues.

### Statistical analysis

All experiments were performed in triplicates, and each data was repeated at least three times except for the epitope mapping experiments, which were performed in duplicates. Graphs are plotted using Graphpad Prism 10.1.2 and represented as mean standard deviations (n = 3). Results were compared by performing a two-tailed Student’s t-test. Significance is defined as P < 0.05, and statistical significance is indicated with an asterisk (*). *P < 0.001 were considered statistically significant.

## Acknowledgements

Authors sincerely acknowledge the contribution of the volunteers participating in this study. The authors also acknowledge staffs of the medical institutions who were involved in this study for support in blood sample collection and handling.

## Funding

This work is primarily supported by a grant from Indian Council for Medical Research (VIR/COVID-19/19/2021/ECD-I) awarded to A.M. and another grant from Bill and Melinda Gates Foundation (INV-042471) awarded to RV. AM also acknowledges the Science and Engineering Research Board (SERB-DST (CRG/2022/003628) for providing additional financial support. R.V. is a JC Bose Fellow of DST. RV also acknowledges infrastructural support was from DST FIST, MHRD, and the DBT IISc Partnership Program. I.D.J. acknowledges the Science and Engineering Research Board (SERB) for the National Post Doctoral Fellowship ([PDF/2023/000284](https://serbonline.in/SERB/ProposalInfoNew)). K.K. acknowledges the support of an MHRD-IISc doctoral fellowship. The funders had no role in study design, data collection and interpretation, or the decision to submit the work for publication.

## Authors’ Contribution

Conceptualization—A.M., and R.V. Designing experiments—A.M., R.V., I.D.J. and K.K. Performing experiments—I.D.J., K.K., and S.R. Manuscript writing—A.M., R.V., I.D.J. and K.K. Fund acquisition—A.M., R.V., B.M., and S.C. Manuscript editing— A.M., R.V., I.D.J. and K.K., B.M., and A.C. Deep Sequencing reads processing—M.B. Medical supervisors in cohort identification and blood collection—S.J., A.C., and I.B.

## Declaration of interests

The authors declare no competing financial interests or dispute in the relationships that could have appeared to influence the work reported in this article.

---

FIG S1. Neutralization of the WA.1, B.1.617.2 and BA.1 pseudoviruses with nine serum samples collected from the selected volunteers during 1st batch (early) (A) and second batch (late) (B) of sample collection. 100% neutralization is considered as obtained by the Polyclonal SARS CoV-2 (NB100-56578) antibody. All data shown are averages of the results of at least three independent experiments ± SD.

Fig S2. (A) The binding profile of the selected serum samples to wild type RBD displayed on the yeast surface. (B-D) Histograms showing yeast cells binding to the indicated serum samples, were sorted into eight gates based upon low to high binding affinity and the frequency of each variant’s barcode in individual gates was determined by Illumina sequencing.

**Fig S3.** Validation of the epitope mapping results was carried out by quantifying the binding of WT and the selected RBD mutants to BCRTH-01, INK09 and INK36 serum (A-C) Histograms showing binding of the yeast cells displaying WT or mutant RBDs to the indicated serum samples, sorted into gates based upon low to high binding. Each image is a representative of two independent experiments performed in triplicates.

**FIG S4.** Multiple sequence alignment of different sarbecovirus spike RBD sequences showing conservation of the Class5/ RBD epitope residues marked by black triangles.

**FIG S5.** Shortlisted 29 sera show neutralising activity against XBB. 1.5. Neutralisation curves for WA. 1 and XBB. 1.5 pseudoviruses with serial dilutions of all the 29 shortlisted sera as indicated.

## Acknowledgement

This study is supported by the Indian Council for Medical Research (VIR/COVID-19/19/2021/ECD-I) awarded to AM. A.M. acknowledges additional funding support from the Anusandhan National Research Foundation (SERB-DST (CRG/2022/003628). R.V. is a JC Bose Fellow of DST. This work was funded by a grant from the Bill and Melinda Gates Foundation (INV-042471) to RV. Funding for infrastructural support was from DST FIST, MHRD, and the DBT IISc Partnership Program. I.D.J. acknowledges the Science and Engineering Research Board (SERB) for the National Post Doctoral Fellowship (PDF/2023/000284) awarded to her. K.K. acknowledges the support of an MHRD-IISc doctoral fellowship. The authors acknowledge project staff Shyamal Manna for his support in blood collection. The funders had no role in study design, data collection and interpretation, or the decision to submit the work for publication.

## Notes

### Competing Interest Statement

The authors have declared no competing interest.

### Summary of Updates

The acknowledgement section has been revised to include funding details.

